# Host selection in European ectomycorrhizal Basidiomycetes affects their estimated spore volumes

**DOI:** 10.1101/2024.07.22.604628

**Authors:** Hironari Izumi

## Abstract

Many ectomycorrhizal fungi show selective associations with certain host tree taxa. Their significance has been demonstrated in the vegetative growth of the fungi, such as structuring the fungal community of the mycorrhizal roots. On the other hand, our understanding of the effects of host selection on reproductive traits such as pore production is very limited. Using two independently published descriptions of host associations and spore dimensions, the differences in spore volumes depending on the associated host taxa were examined in species belonging to 10 ectomycorrhizal genera. The species in *Suillus* were the most host-selective while those in *Inocybe* were generalist. Some species in *Suillus, Lactarius, Russula*, and *Hebeloma* produced significantly (P<0.05) larger spores when they associated with certain host taxa or groups, including Pinaceae or broad-leaved hosts. However, such differences were not found in *Leccinum* and *Tricholoma*. The ecological/evolutionary processes to facilitate host selection through spore volumes were discussed.

## Introduction

Ectomycorrhizas are a symbiotic lifestyle of the soil fungi with woody plant hosts, including Pinaceae, Fagaceae, Betulaceae, and Salicaceae (Alexander, 2006). The recent study reporting that more than half of the ectomycorrhizal fungi are associated with two or fewer host plant genera (Voller et al., 2023) suggests many ectomycorrhizal fungi are selective in their host plant taxa. The host selection of the ectomycorrhizal fungi plays a pivotal role in structuring the fungi’s belowground community (Ishida et al., 2007; Matsuoka et al., 2020). Additionally, the host specificity has been demonstrated to contribute to the dominance of the specialised fungi in the soil fungal community (Tedersoo et al., 2024) by providing putative competitive advantages to the fungi (Kennedy et al., 2020). Therefore, the importance of host selection of ectomycorrhizal fungi on their belowground vegetative growth is well recognised whereas its influence on aboveground reproductive traits, such as sporocarp and spore productions, is still poorly understood, despite the fact that some of the reproductive traits of the ectomycorrhizal fungi were influenced by the host tree. For instance, the host tree sizes affect sporocarp production of associated ectomycorrhizal fungi (Nara et al., 2003) and symbiotic ectomycorrhizal fungi yield larger sporocarps compared to solitary saprotrophs through the possible supply of resources from the hosts (Bässler et al., 2015). Therefore, it is likely that some other reproductive traits of the ectomycorrhizal fungi, such as spore production, are affected by the associated host taxa.

In this study, I tested the possibility that the spore volumes show any differences depending on the host taxa with which they are associated. Ten ectomycorrhizal genera, including *Amanita, Cortinarius, Hebeloma, Hygrophorus, Inocybe, Lactarius, Leccinum, Russula, Suillus, Tricholoma*, were selected for the present study because they are proven to be ectomycorrhizal (Comandini et al., 2012), species-rich, show some host specificity and produce large sporocarps with measurable spores. Additionally, estimated sporocarp biomass of the species was compared between different host taxa to assess the host effects on whole reproductive organ.

## Materials and Methods

Descriptions of the associating host taxa and two-dimensional (length and width) spore size measurements by species were extracted from the books by Læssøe and Petersen (2019) and by Kibby (2020–2023). More than 500 (Kibby, 2020–2023) and 300 (Læssøe and Petersen, 2019) species descriptions were used for the estimation of the spore volumes (Supplemental table 1).

To obtain a sufficient number of species for statistical analysis to evaluate the host effects, the host tree taxa were classified by the family level for *Cortinarius, Hygrophorus, Lactarius, Leccinum, Russula* and *Tricholoma*. For less selective fungi, such as *Amanita, Hebeloma*, and *Inocybe*, the hosts were grouped into broad-leaved only or both broad-leaved and coniferous species. For highly specific genus of *Suillus*, the hosts were divided into genus level, such as *Larix* and *Pinus*. The comparison was made between two major host tree taxa or groups within each ectomycorrhizal genus. To estimate a degree of host specificity in the genus of interest, the proportions of the species associated with a single genus of the host tree were calculated.

The midpoints of length and width measurements were employed to calculate the spore volume for each species (Bässler et al., 2015), using the formula by Calhim et al. (2018). For the volume estimation, the shape of the spores was assumed to be an ellipsoid (Bässler et al., 2015), except for some species of *Amanita* whose shapes were spherical. In that case, the volume was calculated by the formula of V=4/3*3.14*r^3^.

For the comparisons of sporocarp biomass, the measurements were extracted from the descriptions by Kibby only because Læssøe and Petersen did not provide such information. The proxy values of sporocarp biomass were obtained by the squared mean diameter of the cap (Bässler et al., 2015), Brunner-Menzel test was performed to make statistical comparisons using Jamovi software package (The Jamovi project, 2022).

## Results

Among the fungal genera examined, *Suillus* was the most specific genus and all species were associated with a single host tree genus. On the other hand, the least specific i.e., generalist, was *Inocybe* with less than 20% of a single genus associated species (Fig. 1). The spore volumes of the species belong to *Suillus* were significantly (P<0.05) different, depending on the host genera in both descriptions by Kibby and by Læssøe &Petersen. However, those of *Inocybe* were statistically indistinguishable between the hosts by either description. *Suillus, Russula, Lactarius* and *Hebeloma* were the genera that statistically significant differences of spore volumes in specific host taxa were supported by both descriptions. A general trend that the high proportions of the species selecting specific host taxa accompanied with the robustness (supported by two descriptions) of host dependent spore volume difference was observed.

**Fig. 1.**
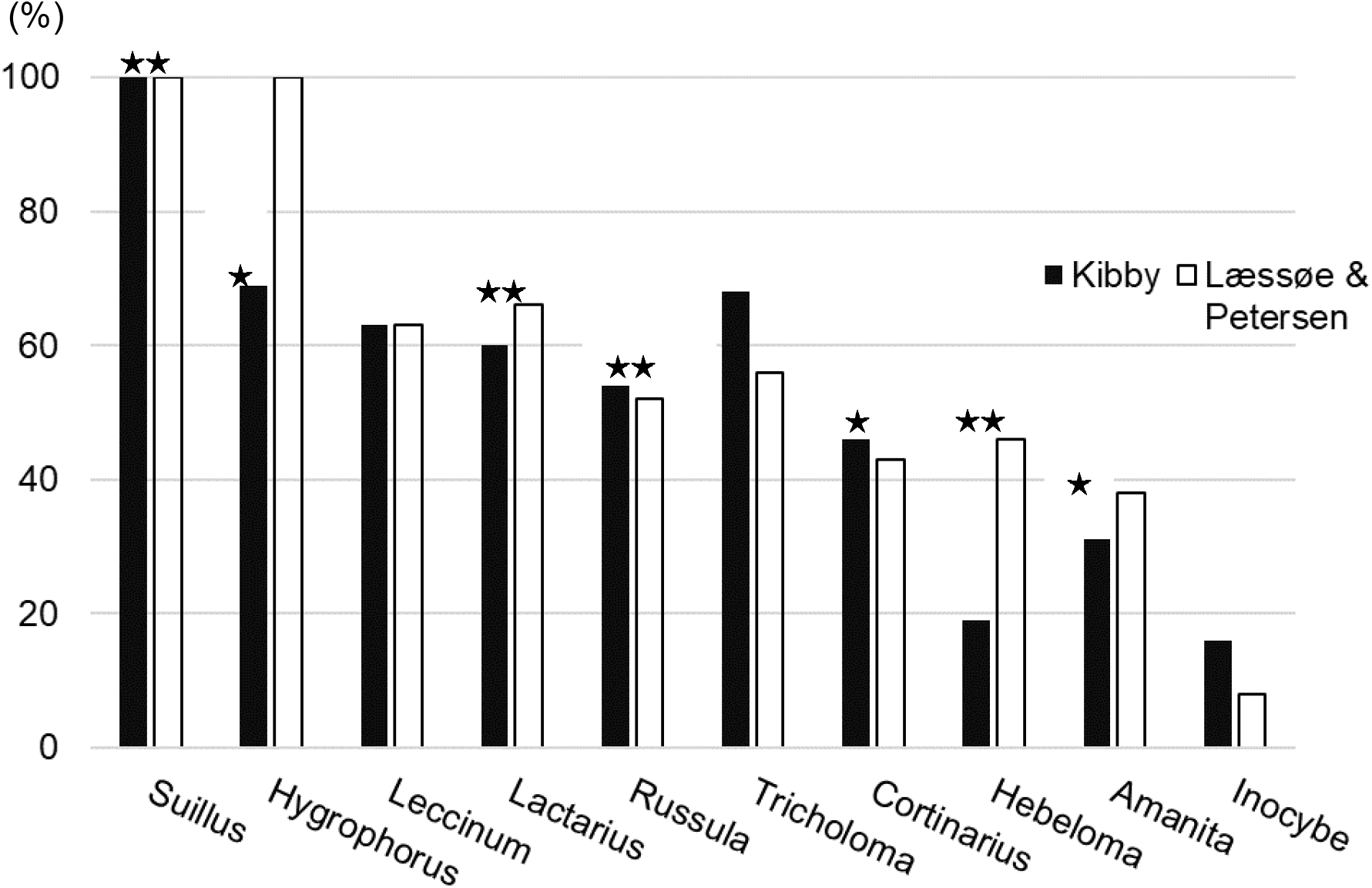
Proportions of the fungal species occurring with a single genus of the host tree. Two independent descriptions of the species by Kibby (2020–2023) and Læssøe and Petersen (2019) are shown in closed and open bars respectively. The star on the bar indicates the fungal descriptions that statistically supported (P<0.05) spore volume differences are observed between species occurring in different host taxa.

In *Russula* and *Lactarius*, the spore volumes of the species associated with Pinaceae hosts were significantly (P<0.05) larger (1.3 folds) than those with Fagaceae or Betulaceae hosts in both descriptions (Table 1). Additionally, the species of *Hebeloma* associated with broad-leaved hosts and those of *Suillus* with *Larix* produced significantly larger (1.3 folds) spores than those with other hosts, such as both of broad-leaved and coniferous or *Pinus* respectively (Table 1). Moreover, some species in *Amanita, Cortinarius* and *Hygrophorus* produced larger spores (1.5 folds, 1.4 folds and 1.3 folds respectively) when they associated with Pinaceae, Fagaceae or broadleaved hosts respectively although statistical significance was observed in only one of two descriptions (Table 1).

**Table 1.**
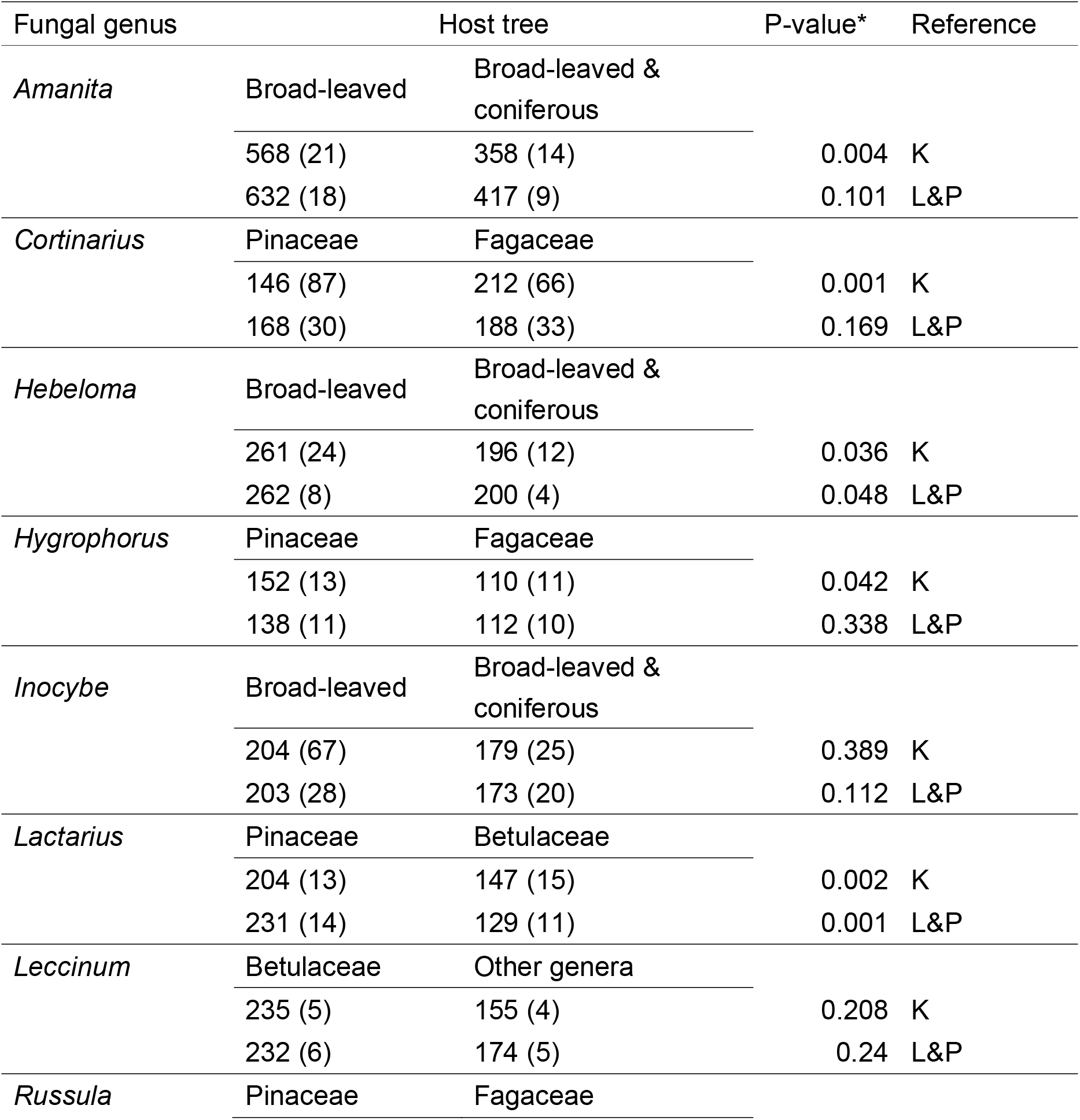
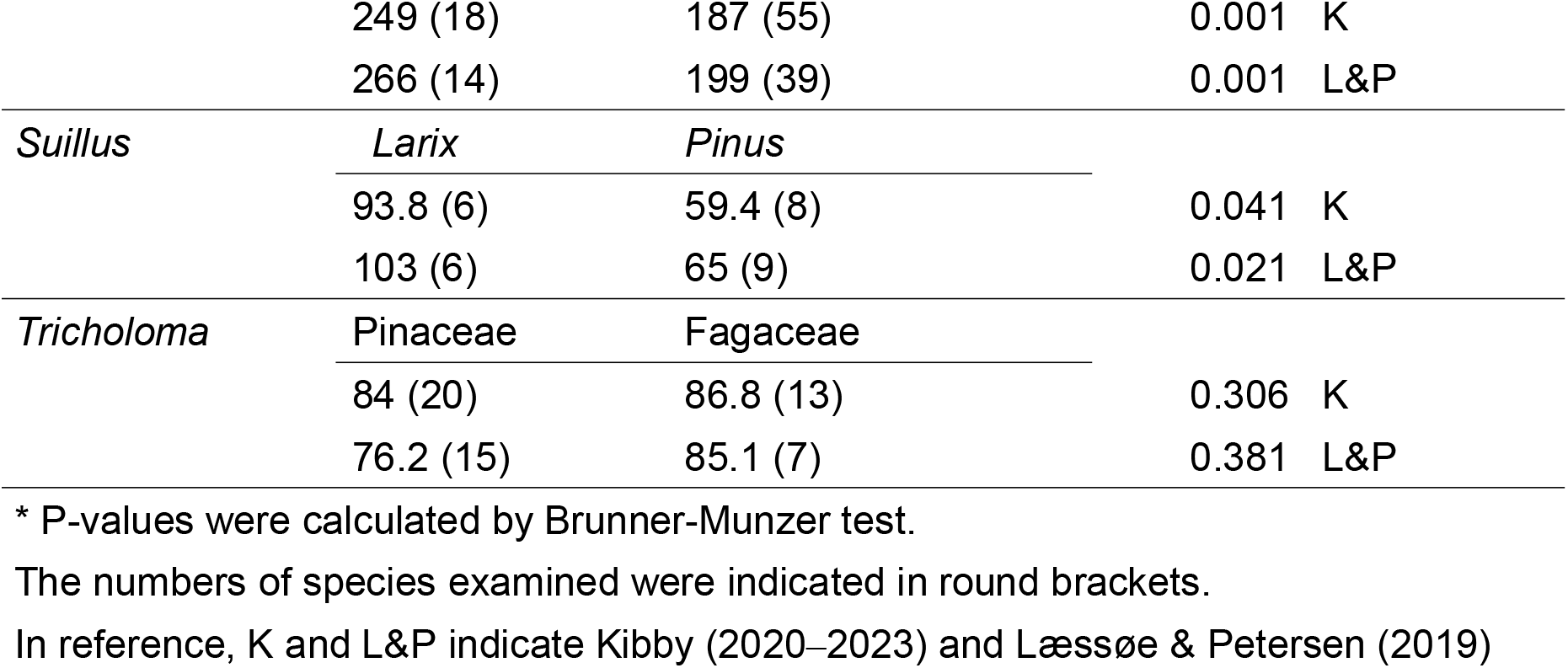
Comparisons of the estimated mean spore volumes (µm^3^) of 10 ectomycorrhizal fungal genera occurred between two different host taxa or groups.

To examine the possibility that observed larger spores in species of *Hebeloma, Lactarius, Russula* and *Suillus* in association with the certain hosts is due to the production of larger sporocarp, the sporocarp biomass was estimated for the same species. There was no statistically significant difference of sporocarp biomass between different host three taxa (Table 2), suggesting the host effect on spore volume is specific.

**Table 2.** Differences of estimated sporocarp sizes occurring between different host tree.

| Fungal genus | Compared host tree combination | P-value |
| --- | --- | --- |
| *Hebeloma* | Broad-leaved/Broad-leaved & Coniferous | 0.209 |
| *Lactarius* | Pinaceae/Betulaceae | 0.944 |
| *Russula* | Pinaceae/Fagaceae | 0.651 |
| *Suillus* | Larix/Pine | 0.548 |

## Discussion

Because the fungi forming ectomycorrhizas with plants are diverse and consist of, at least, 5400 species (Sharma, 2017), the number of species examined in this study is rather small (500 species at most). Although such a small number would provide a partial picture of the host effects on ectomycorrhizas interesting patterns emerge. Firstly, degrees of host selection vary depending on the fungal taxa. Some species belonging to the genera *Hebeloma, Lactarius, Russula*, and *Suillus* tend to be more host-selective than those in Inocybe (Fig, 1). Secondary, they produce larger spores in association with certain host taxa (Table 1). The host selectivity observed in this study can arise from two distinct pathways. One is phynotypic flexibility of the fungi to the current environmental conditions. The other is fungal genotypic evolution to increase fitness by associating with particular host taxa. It is well known that overall sporocarp production is influenced by rainfall and ambient temperature (Boddy et al., 2014). Thus, there is a possibility that observed larger spores in association with certain hosts may reflect phynotipic adaptation to facilitate survival under the conditions which the host grows.

However, phynotypic adaptation may not play critical role on the spore volume difference because the result in this study showing no statistically significant difference of sporocarp biomass between different suggests that the sporocarp sizes of the species are consistent within the genus and it is likely that spore volumes are also genetically fixed in the same genus. These lines of reasoning lead us to conclude that evolutionally adaptation is more plausible for developing host specificity with larger spore volumes. Then, what is the advantages to evolve producing larger spores? As the spores are reproductive organ the advantages must be placed on how much the larger spore volumes contribute fungal dispersal and survival. Indeed, some tropical *Amanita* produce smaller spores for effective areal dispersal between sparsely distributed hosts (Tulloss, 2005). The larger spores, on the other hand, would have merits for survival rather than dispersal. In this study, *Larix* spp., the host of highly specific *Suillus*, grows under dry, cold winter conditions in native high Alpine rages and usually performs better in poor to medium nutrient sites as an early coloniser in the lower elevation (Da Ronch et al., 2016). Additionally, it has been shown that the larger spores were thought to retain more water and provide some protection against desiccation (Kauserud et al., 2011) and spores become larger in resource-limited habitats (Halbwachs et al., 2017).

Therefore, larger spores in *Larix*-specific *Suillus* species may be the evolutionary consequences selected by better survival of the larger spores in harsh environments. *Russula, Lactarius* and *Hebeloma* have been known to be “late-stage” fungi (Smith & Read, 2008; Ishida et al., 2008). The sporocarps of *Lactarius*, for instance, appear at older stand ages in *Pinus sylvestris* forests (Bonet et al., 2004) and those of *Russula* and *Hebeloma* are observed in a late stage of primary vegetative succession on volcanic Mt Fuji (Nara et al., 2003). Whereas “late-stage” fruiting may be due to slow developments of belowground fungal biomass, the late fruiting may be caused by delayed spore germination, waiting for the time when the host tree reaches a certain age. The previous report showing that the spores of these genera are difficult to produce mycorrhiza by inoculations of the spores to seedlings (Nara, 2009) suggests spore germination may require more mature host trees. Thus, larger spores can contribute to remaining viable for a long time by waiting for the host with suitable ages to be around. This also explains why *Inocybe* does not show host-specific differences in spore volumes. Inocybe has been suggested one of early colonisers in primary vegetative succession (Nara et al., 2003), which they can immediately colonise the host and do not wait for colonisation for long.

How the spores become larger when the fungi are associated with certain host taxa is completely unexplored but one possibility is that specific association may increase physiological affinity between the fungi and the host, which facilitates to produce the larger spores. The previous study has demonstrated that host tree allocated more carbon to the specific *Suillus grevillei* colonised roots compared to non-specific *S. bovinus* (Finlay,1989). Therefore, it is likely that host-specific fungi obtain more photo-assimilated carbons from specific hosts and can produce larger spores. The interesting genera are *Leccinum* and *Tricholoma*, which have high (>50%) host-selective species but no statistically significant host effects on the spore volume. *Leccinum* has been considered to shift their hosts recently (den Bakker et al., 2004), which may not provide sufficient time to develop the host effects on spore volumes. *Tricholoma* spp. have been shown to form mycorrhizas between oak and pine in vitro (Yamanaka et al., 2014). Thus, the observed selectivity in the field descriptions may reflect the environmental conditions, such as the availability of the host trees in the locations the fungi grow. These two instances, of *Leccinum* and *Tricholoma*, suggests that stable and exclusive association between the host and the fungi are required to manifest the host effects on the spore volume.

The spore volumes of *Amanita, Cortinarius*, and *Hygrophorus* increased in association with the hosts of broad-leaved or Pinaceae respectively (Table 1) alth ough only one of the two descriptions, namely by Kiddy, was statistically supported. As Kiddy describes mainly British species, this may reflect more specialisation to the host taxa of the species in these genera in Britain, by geographical isolation from mainland Europe. For example, the geographical isolation by growing on an island, such as Corsica, can change the relationship between ectomycorrhizal fungi and their host tree by shifting the host ranges of the fungi (Rochet et al., 2011). Thus, it is possible that the separation of British species from those in mainland Europe may also influence the host effects on spore volumes although this requires more detailed examinations.

## Disclosure

The author declares no conflicts of interest.

## Acknowledgment

This work was financially supported by the Jurinji Buddhist temple. Dr Melanie Roy of PCI Ecology was acknowledged for her feedback and encouragement for early version of the manuscript.

## Notes

### Competing Interest Statement

The authors have declared no competing interest.

### Summary of Updates

The discussion section of the manuscript was reworked for improvements.

